# The genetics of an early Neolithic pastoralist from the Zagros, Iran

**DOI:** 10.1101/059568

**Authors:** M. Gallego-Llorente, S. Connell, E. R. Jones, D. C. Merrett, Y. Jeon, A. Eriksson, V. Siska, C. Gamba, C. Meiklejohn, R. Beyer, S. Jeon, Y. S. Cho, M. Hofreiter, J. Bhak, A. Manica, R. Pinhasi

**Affiliations:** Department of Zoology, University of Cambridge, Cambridge,CB2 3EJ, UK.; School of Archaeology and Earth Institute, University College Dublin, Belfield, Dublin 4, Ireland.; Department of Archaeology, Simon Fraser University, Burnaby, BC, Canada, V5A 1S6.; The Genomics Institute, Ulsan National Institute of Science and Technology (UNIST), Ulsan 44919, Republic of Korea.; Department of Biomedical Engineering, School of Life Sciences, Ulsan National Institute of Science and Technology (UNIST), Ulsan 44919, Republic of Korea.; Integrative Systems Biology Laboratory, Division of Biological and Environmental Sciences & Engineering, King Abdullah University of Science and Technology, Thuwal 23955-6900, Kingdom of Saudi Arabia.; Smurfit Institute of Genetics, Trinity College Dublin, Dublin, Dublin 2, Ireland.; Centre for GeoGenetics, Natural History Museum of Denmark, University of Copenhagen, Øster Voldgade 5-7, Copenhagen 1350, Denmark.; Department of Anthropology, University of Winnipeg, Winnipeg, MB, Canada, R3B 2E9.; Evolutionary Adaptive Genomics, Institute for Biochemistry and Biology, Department of Mathematics and Natural Sciences, University of Potsdam, Karl-Liebknechtstraße 24-25, 14476 Potsdam, Germany.

## Abstract

The agricultural transition profoundly changed human societies. We sequenced and analysed the first genome (1.39×) of an early Neolithic woman from Ganj Dareh, in the Zagros Mountains of Iran, a site with early evidence for an economy based on goat herding,ca. 10,000 BP. We show that Western Iran was inhabited by a population genetically most similar to hunter-gatherers from the Caucasus, but distinct from the Neolithic Anatolian people who later brought food production into Europe. The inhabitants of Ganj Dareh made little direct genetic contribution to modern European populations, suggesting they were somewhat isolated from other populations in the region. Runs of homozygosity are of a similar length to those from Neolithic Anatolians, and shorter than those of Caucasus and Western Hunter-Gatherers, suggesting that the inhabitants of Ganj Dareh did not undergo the large population bottleneck suffered by their northern neighbours. While some degree of cultural diffusion between Anatolia, Western Iran and other neighbouring regions is possible, the genetic dissimilarity of early Anatolian farmers and the inhabitants of Ganj Dareh supports a model in which Neolithic societies in these areas were distinct.

## Introduction

The agricultural transition started in a region comprising the Ancient Near East and Anatolia ~12,000 years ago with the first Pre-Pottery Neolithic villages and the first domestication of cereals and legumes (*1*, *2*). Archaeological evidence suggests a complex scenario of multiple domestications in a number of areas (*3*), coupled with examples of trade (*4*). Ancient DNA (aDNA) has revealed that this cultural package was later brought into Europe by dispersing farmers from Anatolia (the so called ‘demic’ diffusion, as opposed to non-demic cultural diffusion (*5*–*10*)) ~8,400 years ago. However a lack of aDNA from early Neolithic individuals from the Near East leaves a key question unanswered: was the agricultural transition developed by one major population group spanning the Near East, including Anatolia; or was the region inhabited by genetically diverse populations, as is suggested by the heterogeneous mode and timing of the appearance of early domesticates at different localities? Based on the archaeological record there is a major difference between populations in the Levant, Mesopotamia and Anatolia, whose early Neolithic subsistence is based on cereals and legumes (classical agriculturalists) and those from the Zagros, who began the herding of goats and sheep and for whom there is little evidence of true cultivation (classical pastoralists).

To answer this question, we sequenced the genome of an early Neolithic female from Ganj Dareh, GD13a, from the Central Zagros (Western Iran), dated to 10000)9700 cal BP (*11*), a region located at the eastern edge of the Near East. Ganj Dareh is well known for providing the earliest evidence of herd management of goats beginning at 9,900 BP (*11*–*13*). It is a classic mound site at an altitude of ~1400m in the Gamas)Ab Valley of the High Zagros zone in Kermanshah Province, Western Iran. It was discovered in the 1960s during survey work and excavated over four seasons between 1967 and 1974. The mound, ~40 m in diameter, shows 7 to 8 m of early Neolithic cultural deposits. Five major levels were found, from top to bottom A through E. Extended evidence showed a warren of rooms with evidence of burial under floors and within what may be burial chambers and/or disused houses (*14*). The current MNI is 116, with 56 catalogued as skeletons and having four or more bones recovered (*15*). The individual analysed was part of burial 13, containing three individuals and recovered in level C in 1971 from the floor of a brick-walled structure. The individual sampled, 13A, was a 30-50 year old female; the other individuals in the burial unit were a second adult (13B) and an adolescent (13).

The site has been directly dated to 9650-9950 calBP (*11*), showing intense occupation over two to three centuries. The economy of the population has been shown to be that of pastoralists, focusing on goats (*11*). Archaeobotanical evidence is limited (*16*) but the evidence present is for two-row barley, probably wild, and no evidence for wheat, rye or other domesticates. In other words the overall economy is divergent from the classic agricultural mode of cereal agriculture found in the Levant, Anatolia and Northern Mesopotamian basin.

### Results

The petrous bone of GD13a yielded sequencing libraries comprising 18.57% alignable human reads that were used to generate 1.39-fold genome coverage. The sequence data showed read lengths and nucleotide misincorporation patterns indicative of post)mortem damage, and had low contamination estimates (<1%, see Supplementary).

We compared GD13a with a number of other ancient genomes and modern populations (*6*, *17*–*29*), using principal component analysis (PCA) (*30*), ADMIXTURE (*31*) and outgroup *f*_3_ statistics (*32*) (Fig. 1). GD13a did not cluster with any other early Neolithic individual from Eurasia in any of the analyses. ADMIXTURE and outgroup *f*_3_ identified Caucasus Hunter-Gatherers of Western Georgia, just north of the Zagros mountains, as the group genetically most similar to GD13a (Fig. 1B&C), whilst PCA also revealed some affinity with modern Central South Asian populations such as Balochi, Makrani and Brahui (Fig. 1A and Fig. S4). Also genetically close to GD13a were ancient samples from Steppe populations (Yamanya & Afanasievo) that were part of one or more Bronze age migrations into Europe, as well as early Bronze age cultures in that continent (Corded Ware) (*17*, *23*), in line with previous relationships observed for the Caucasus Hunte-Gatherers (*26*).

**Fig. 1.**
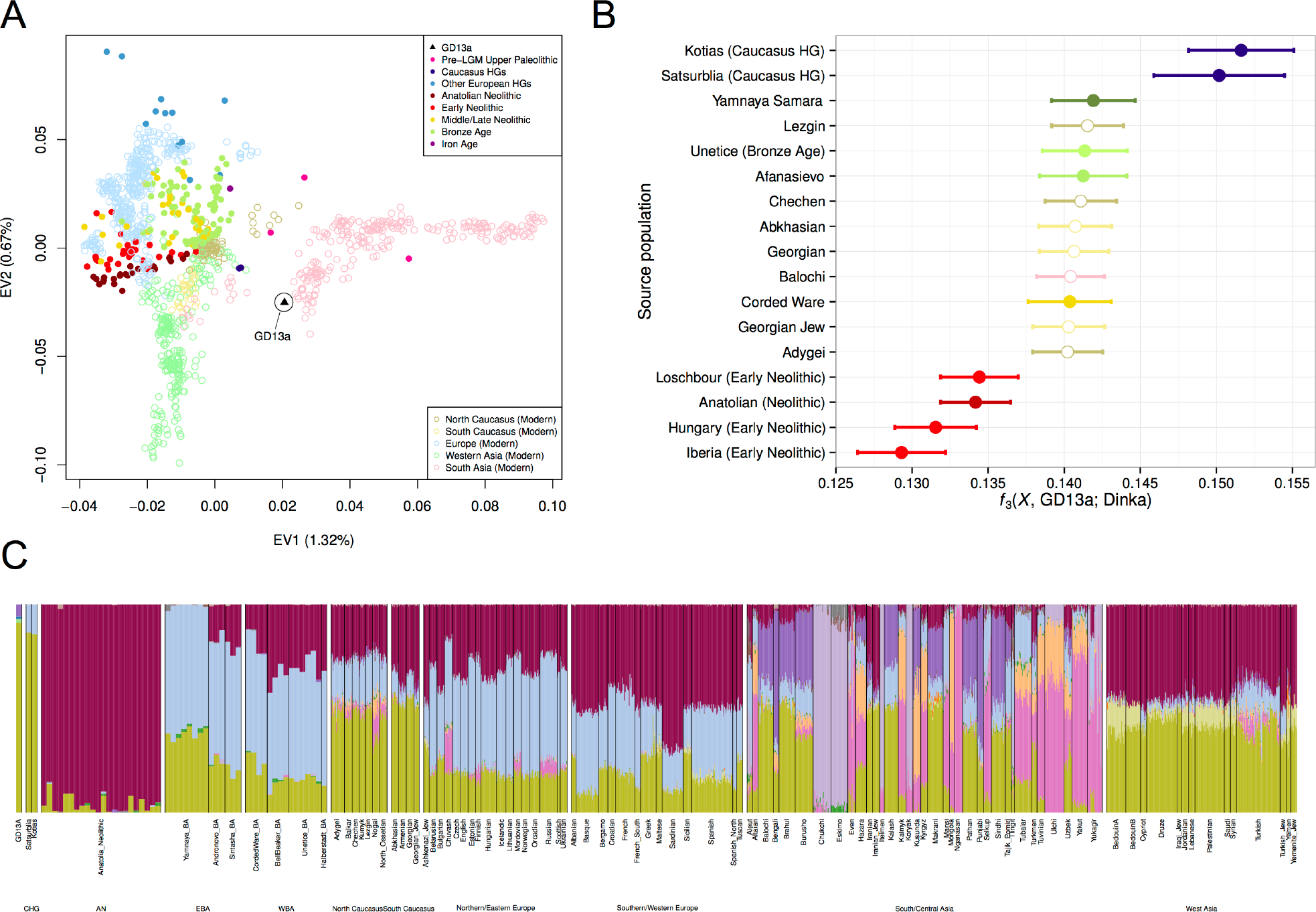
GD13a appears to be related to Caucasus Hunter Gatherers and to modern South Asian populations. **A) PCA** loaded on modern populations (represented by open symbols). Ancient individuals (solid symbols) are projected onto these axes. **B) Outgroup *f*_3_(*X*, GD13a; Dinka)**, where Caucasus Hunter Gatherers (Kotias and Satsurblia) share the most drift with GD13a. Ancient samples have filled circles whereas modern populations are represented by empty symbols. **C) ADMIXTURE** using K=17, where GD13a appears very similar to Caucasus Hunter Gatherers, and to a lesser extent to modern south Asian populations.

We further investigated the relationship between GD13a and Caucasus Hunter-Gatherers using *D*-statistics (*32*, *33*) to test whether they formed a clade to the exclusion of other ancient and modern samples. A large number of Western Eurasian samples (both modern and ancient) showed significant excess genetic affinity to the Caucasus Hunter-Gatherers, whilst none did with GD13a. Overall, these results point to GD13a having little direct genetic input into later European populations compared to its northern neighbours.

To better understand the history of the population to which GD13a belonged, we investigated the distribution of lengths of Runs of Homozygosity (ROH) (Fig. 2a). A bias towards a high frequency of long ROH is indicative of past strong bottlenecks, whilst a high frequency of shorter ROH suggests a relatively large stable population. GD13a has a distribution with more short ROH (<2 Mb), similar to that of the descendant of Anatolian early farmers (represented by the European farmers NE1 (*21*) and Stuttgart (*18*)). In contrast, both Western (*18*) and Caucasus Hunter-Gatherers (*26*) both have relatively more long ROH (>2 Mb). Thus, GD13a is the descendant of a group that had relatively stable demography and did not suffer the bottlenecks that affected more northern populations.

**Fig. 2.**
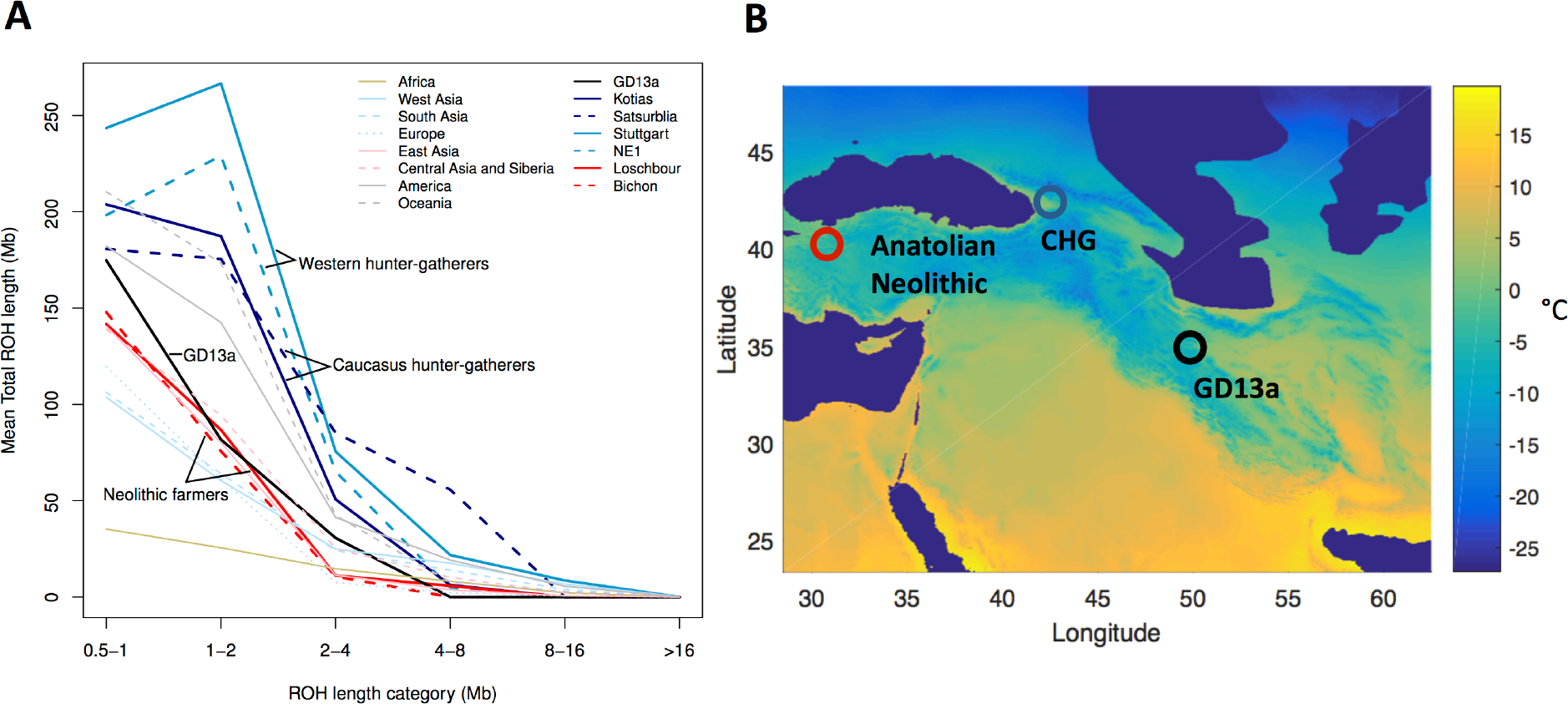
GD13a did not undergo a large population bottleneck during the LGM. **A)** GD13a has similar runs of homozygosity (ROH) lengths to Neolithic individuals, while Caucasus Hunter Gatherers (Kotias and Satsurblia), like European Hunter Gatherers (Loschbour and Bichon), underwent large population bottlenecks during the LGM. **B)** Map showing geographical location of Anatolian Neolithic samples, Caucasus Hunter Gatherers (CHG) and GD13a. Background colours indicate mean temperature (°C) of coldest quarter during the LGM, with LGM sea levels. **Map of populations was generated with R software** (v3.1.2, https://cran.r-proiect.org/) (*52*)

The phenotypic attributes of GD13a are similar to the neighbouring Anatolian early farmers and Caucasus Hunte-Gatherers. Based on diagnostic SNPs, she had dark, black hair and brown eyes (see Supplementary). She lacked the derived variant (rs16891982) of the *SLC45A2* gene associated with light skin pigmentation but had at least one copy of the derived *SLC24A5* allele (rs1426654) associated with the same trait. The derived *SLC24A5* variant has been found in both Neolithic farmer and Caucasus hunter)gatherer groups (*5*, *21*, *26*)suggesting that it was already at appreciable frequency before these populations diverged. Finally, she did not have the most common European variant of the *LCT* gene (rs4988235) associated with the ability to digest raw milk, consistent with the later emergence of this adaptation (*5*, *21*, *23*).

It is possible that farmers related to GD13a contributed to the eastern diffusion of agriculture from the Near East that reached Turkmenistan (*34*) by the 6^th^ millennium BP, and continued further east to the Indus Valley (*35*). However, detecting such a contribution is complicated by a later influx from Steppe populations with Caucasus Hunter-Gatherer ancestry during the Bronze Age. We tested whether the Western Eurasian component found in Indian populations can be better attributed to either of these two sources, GD13a and Kotias (a Caucasus Hunter Gatherer), using D-statistics to detect gene flow into an ancestral Indian component (represented by the Onge). For all tests where a difference could be detected, Kotias acted a better proxy than GD13a (Fig. S9 and Table S6). This result implies that the majority of the West Eurasian component seen in India derives from the Bronze age migrations; this interpretation is supported by dating of last contact based on patterns of Linkage Disequilibrium (*36*).

### Discussion

GD13a had little direct genetic input into later European populations compared to the Caucasus Hunter-Gatherers (its northern neighbours) as demonstrated using D-statistics. This lack of connectivity with neighbouring regions might have arisen early on, since we also find that Hunter-Gatherers from the Caucasus show excess affinity to Western Hunte-Gatherers and early Anatolian farmers; this result suggests the possibility of gene flow between the former and these two latter groups to the exclusion of GD13a. An alternative, but not mutually exclusive, explanation for this pattern is that GD13a might have received genetic input from a source equally distant from all other European populations, and thus basal to them.

The Last Glacial Maximum (LGM) made entire regions in northern Eurasia uninhabitable, and therefore a number of hunter-gatherer populations likely moved to the south. For Europe there may be a separation between Western and Eastern populations with minimal occupation of the Central European plains (*24*). For Eastern Europe, Central Asia and the northern Near East, glaciation itself was less a factor (*26*). In these areas, our understanding of how populations weathered the LGM is still vague at best. It has previously been suggested that differences in the frequency of long and short runs of homozygosity in ancient samples may be associated with different demographic experiences during the LGM (*36*). Anatolian farmers, with their shorter ROH, have been argued to have been relatively little affected by the LGM compared to Western and Caucasus Hunter) Gatherers, which are characterised by more long ROH (>2Mb). GD13a has a profile similar to that of Anatolian, suggesting that her ancestors also faced more benign conditions compared to populations further north. Superimposing the sampling locations of these groups onto climatic reconstructions from the LGM (Fig. 2b), however, does not reveal clear climatic differences among the regions. It is possible that the ancestors of the Anatolian farmers and Ganj Dareh spent the LGM in areas further south or east, which experienced milder climate. But it is also possible that they exploited local pockets of favourable climate. Whilst high elevation sites in the Zagros were abandoned during the LGM(*37*), there are a number of sites in the valleys that were occupied during that period and might have experienced more favourable conditions.

The archaeological record indicates an eastward Neolithic expansion from the eastern regions of the Near East into Central and South Asia (*34*, *38*). Our analysis shows that the Caucasus Hunter Gatherer Kotias acts as a better proxy than GD13a for the Eurasian Ancestry found in that part of Asia. However, given the similarity between the two sources, the possibility remains that an older, smaller contribution from a source genetically close to GD13a would be masked by later gene flow from the Steppe. Eventually, ancient DNA from the Indus Valley will be needed to detect conclusively whether any genetic traces were left by the eastward Neolithic expansion from the Near East, or whether this process was mostly cultural.

The presence of two distinct lineages in the Near East at the beginning of the Neolithic transition raises an interesting question regarding the independence of innovations arising at different locations. Ganj Dareh has the earliest known evidence for goat domestication (*11*–*13*), and the Zagros mountains have also been argued to have the site of early farming (*3*). Were these innovations independent of similar achievements that made up the Neolithic package that North West Anatolians brought into Europe? Or were they exchanged culturally? If the latter, it would imply a cultural diffusion in the absence of much genetic interchange.

## Methods

### DNA extraction and library preparation

Sample preparation, DNA extraction and library preparation were carried out in dedicated ancient DNA facilities at University College Dublin. The dense part of the petrous bone was isolated, cleaned and sequenced following experimental procedures outlined in. DNA was extracted from 310 mg of ground bone powder using a double-digestion and silica column method as described in. Indexed Illumina sequencing libraries were constructed with a protocol based on with modifications including blunt end repair using NEBNext End Repair Module (New England BioLabs Inc) and heat inactivation of Bst DNA polymerase.

### Sequence processing and alignment

Libraries were sequenced over a flow cell on a HiSeq 2000 at the Theragen BiO Institute (South Korea) using 100 bp single-end sequencing. Adapter sequences were trimmed from the 3′ ends of sequences using cutadapt version 1.3, conservatively requiring an overlap of 1 base pair (bp) between the adapter and the read. Reads were aligned using BWA, with the seed region disabled, to the GRCh37 build of the human genome with the mitochondrial sequence replaced by the revised Cambridge reference sequence (NCBI accession number NC_012920.1). Data from separate lanes were merged using Picard MergeSamFiles (http://picard.sourceforge.net/) and duplicate reads from the same library amplification were filtered using SAMtools rmdup. Sequences were further filtered to remove those with mapping quality < 30 and length < 30 bp. Indels were realigned using RealignerTargetCreator and IndelRealigner from the Genome Analysis Toolkit. The first and last 2 bp of each read were sofrclipped to a base quality of 2 to reduce the impact of terminal sequence damage on analyses. The average genome-wide depth of coverage was calculated using the *genomecov* function of bedtools. A summary of alignment statistics can be found in Table S1.

### Authenticity of results

The data were assessed for the presence of typical signatures of DNA (*39*, *40*). The sequence length distribution of molecules was examined as outlined in (*41*) (Fig. S2) while the prevalence of nucleotide misincorporation sites at the ends of reads was evaluated using mapDamage 2.0 and a random subsample of 1 million reads (*42*) (Fig. S3).

The mitochondrial contamination rate was assessed by evaluating the proportion of non-consensus bases at haplogroup defining positions in the mitochondrial genome (*21*, *43*). Only bases with quality ≥ 20 were used. The X chromosome contamination rate could not be evaluated as the sample was determined to be female, using the script described in (*44*).

### Mitochondrial Haplogroup Determination

To determine to which haplogroup the mitochondria of GD13a belonged, a consensus sequence was generated using ANGSD (*45*). Called positions were required to have a depth of coverage ≥ 3 and only bases with quality ≥ 20 were considered. The resulting FASTA files were uploaded to HAPLOFIND (*46*) for haplogroup determination. Coverage was calculated using GATK DepthOfCoverage (*47*).

### Dataset preparation for population genetic analyses

Genotypes were called in GD13a at sites which overlapped those in the Human Origins dataset (Lazaridis et al., 2014 (*18*), filtered as described in Jones et al., 2015 (*26*)) using GATK Pileup (*47*). Triallelic SNPs were discarded and bases were required to have quality ≥ 30. For positions with more than one base call, one allele was randomly chosen with a probability equal to the frequency of the base at that position. This allele was duplicated to form a homozygous diploid genotype for each position called in GD13a. This method of SNP calling was also used for selected ancient samples described in (Jones et al., 2015 (*26*) Cassidy et al., 2015 (*27*), Gunther et al. (*28*), Omrak et al., 2015 *6* and Olalde et al., 2015 (*29*)). Genotype calls for these ancient samples were merged with calls from modern samples found in the Human Origins dataset and ancient samples provided in the Mathieson et al. 2005 (*5*) dataset which also included genotype calls for previously published ancient samples (*7*, *26*, *27*, *32*–*37*). To avoid biases caused by post-mortem DNA damage, only transversion sites were used for PCA, ADMIXTURE, *f*_3_-statistics and *D*-statistics.

### Principal component analysis

To explore GD13a and other ancient samples in the context of modern variation in Eurasia, we performed PCA with a panel of contemporary populations (196 contemporary populations, 145,004 transversion SNPs). The analysis was carried out using SmartPCA (30); the components were loaded on the contemporary populations, and the ancient individuals were projected onto these dimensions (Fig. 1 and Fig. S4).

### ADMIXTURE

A clustering analysis was performed using ADMIXTURE version 1.23 (*31*), using the full panel of modern and ancient samples described above. SNPs in linkage disequilibrium were thinned using PLINK (v1.07) (*48*) with parameters -indep-pairwise 200 25 0.5 (*17*), resulting in a set of 116,834 SNPs for analysis. Clusters (K) (2-20) were explored using 3 runs with fivefold cross-validation at each K with different random seeds. The minimal cross-validation error was found at K=17, but the error already starts plateauing from roughly K=10, implying little improvement from this point onwards (Fig. S5). The ADMIXTURE proportions are shown in Fig. S6 for all samples and in Fig. 1c for GD13a and selected modern and ancient populations harbouring the component dominant for GD13a.

### Outgroup *f*_3_-statistics and *D*-statistics

Outgroup *f*_3_-statistics and *D*-statistics were performed using the qp3Pop and qpDstat programs from the ADMIXTOOLS package.

### Neighbour-joining tree

We used a custom Matlab script to calculate pairwise pi from genome-wide genotype data using a panel of 22 individuals (from the dataset described above), including GD13a, representative ancient samples, and different modern populations from the same geographic area as GD13a, and generated a UPGMA tree using the seqlinkage function in Matlab's Bioinformatics Toolbox.(*49*)

### Runs of homozygosity

In order to examine runs of homozygosity (ROH) we used imputation to infer diploid genotypes in our sample following the method described in. We used GATK Unified genotyper to call genotype likelihoods at SNP sites in Phase 3 of 1,000 genomes project and ROH analysis was carried out as outlined in S10.

### Phenotypes of interest

Genes associated with a particular phenotype in modern populations were explored in GD13a. Observed genotypes were called using GATK Unified genotyper (47), calling alleles present in Phase 1 of 1,000 genomes dataset (50) with base quality ≥ 20. As many diagnostic markers had lfold coverage or less, we also used imputation to infer genotypes at these positions. Imputation was performed as described in section S11, imputing at least 1Mb upstream and downstream of the SNP position (this interval was reduced for those variants within the first 1Mb of the chromosome). The Hirisplex prediction model (51) was used to explore hair and eye colour (Table S5). For the observed data, if the sample had 1× coverage, the variant was called as homozygous for that allele. Hair and eye colour predictions were confirmed using imputed data.

## Acknowledgements

A.M. was supported by ERC Consolidator Grant 647787 ‘LocalAdaptation’; R.P. by ERC Starting Grant: ERC– 2010-StG 26344; M.H. by ERC Consolidator Grant 310763 ‘GeneFlow’; C.G. was supported by the Irish Research Council for Humanities and Social Sciences (IRCHSS) ERC Support Programme and the Marie-Curie Intra-European Fellowships (FP7-IEF-328024); J.B. was supported by the 2014 Research fund (1.140077.01) of Ulsan National Institute of Science & Technology (UNIST). and Geromics internal research funding; and M.G. by a BBSRC DTP studentship.

## Author contributions statement

D.C.M, C.M. and R.P. obtained the sample and provided expertise in the archaeology. C.G. and S.C. extracted the genetic material, Y.J, S.J., Y.S.C. and J.B. sequenced the sample, M.G., E.R.J., V.S., A.E. R.B. and A.M. did the genetic analysis. M.H. helped with interpretation.

## Accession codes

Raw reads from Ganj Dareh 13 are available for download through the EBI European Nucleotide Archive (ENA) accession number PRJEB13189.

